# Oxidative stress induces cell death partially by decreasing both mRNA and protein levels of nicotinamide phosphoribosyltransferase in PC12 cells

**DOI:** 10.1101/2020.08.20.259143

**Authors:** Cuiyan Zhou, Weihai Ying

**Affiliations:** Med-X Research Institute and School of Biomedical Engineering, Shanghai Jiao Tong University, Shanghai 200030, P.R. China

**Keywords:** Nampt, NAD^+^, Oxidative Stress, Cell Death, Aging

## Abstract

Numerous studies have indicated critical roles of NAD^+^ deficiency in both aging and multiple major diseases. It is critical to investigate the mechanisms underlying the NAD^+^ deficiency under the pathological conditions. It has been reported that there was a decreased level of Nicotinamide phosphoribosyltransferase (Nampt) – an important enzyme in the salvage pathway of NAD^+^ synthesis – under certain pathological conditions, while the mechanisms underlying the Nampt decrease require investigation. In this study we used differentiated PC12 cells as a cellular model to investigate the effects of oxidative stress on both the mRNA and protein levels of Nampt, as well as the role of this effect in oxidative stress-induced cell death: First, Nampt plays significant roles in both the NAD^+^ synthesis and survival of the cells under basal conditions; second, H_2_O_2_ produced significant decreases in both the mRNA levels and the protein levels of Nampt; and third, H_2_O_2_ induced cell death partially by producing the decreases in the mRNA and protein levels of Nampt, since the Nampt inhibitor or the Nampt activator significantly exacerbated or attenuated the H_2_O_2_-induced cell death, respectively. Collectively, our study has indicated that oxidative stress can decrease both the mRNA and protein levels of Nampt, which has indicated a novel mechanism underlying the NAD^+^ deficiency in aging and under multiple pathological conditions. Our study has also indicated that the decreased Nampt levels contribute to the H_2_O_2_-induced cell death, suggesting a new mechanism underlying oxidative cell death.

## Introduction

Numerous studies have indicated that NAD^+^ plays important roles in mutiple biological functions (1). Increasing evidence has also indicated NAD^+^ deficiency as a critical pathological factor in multiple major diseases and pathological conditions (1, 2, 3), including ischemic brain injury (4, 5), mayocardial ischemia (6), head trauma (7), epilepsy (8), and radiation-induced tissure injury (9) and chemotherapy agent-induced tissue injury (10). NAD^+^ deficiency also appears to play an important role in the aging processess (11–13), which has been the biological basis for the widely used NAD^+^ supplements. It is of critical theoretical and medical significance to expose the mechanisms underlying the pathologocal alteration of NAD^+^ metabolism. However, there has been significant shortage of information on this important topic.

There are two pathways for NAD^+^ synthesis, including the *de novo* pathway and the salvage pathway (14). The nuclear enzyme Nicotinamide mononucleotide adenylyltransferase-1 (NMNAT-1) is a crucial enzyme in both the *de novo* pathway and the salvage pathway of NAD^+^ synthesis (14). Mammals can use nicotinamide as the precursor for NAD^+^ synthesis in the salvage pathway (15): Nicotinamide phosphoribosyltransferase (Nampt) converts nicotinamide to nicotinamide mononucleotide (NMN) that can be converted to NAD^+^ by NMNATs (15).

NAD^+^ deficiency has been found in models of a number of diseases, which may result from poly (ADP-ribose) polymerase-1 (PARP-1) activation and decreased Nampt activity. It has been indicated that decreased Nampt activity is an important mechanism accounting for impaired NAD^+^ synthesis capacity: Decreased Nampt activity has been found in the metabolic organs that are exposed to high-fat diet (HFD) (16), Drosophila pink1 mutants (17), and aging (16, 18). It is critical to investigate the mechanisms underlying the decreased Nampt levels.

In this study we used PC12 cells as a cellular model to investigate the effects of oxidative stress on both the mRNA and protein level of Nampt. We also determined the roles of Nampt in the cell survival and NAD^+^ metabolism in both basal and oxidative stress conditions. Our study has not only indicated significant roles of Nampt in the survival of PC12 cells under both basal and oxidative stress conditions, but also suggested novel mechanisms underlying the decreased Nampt in diseases and aging.

## Materials and methods

### Materials

H_2_O_2_ (323381) was purchased from Sigma Aldrich (St Louis, Missouri, USA). FK866 (HY-50876) and P7C3 (HY-15976) were purchased from MedChemExpress (New Jersey, USA). Nampt siRNAs and control siRNAs were purchased from GenePharma (Shanghai, China). Nampt primers were purchased from Sangon Biotech (Shanghai, China).

### Cell culture

Differentiated PC12 cells were purchased from the Cell Resource Center of Shanghai Institute of Biological Sciences, Chinese Academy of Sciences (Shanghai, China). The cells were plated into 24- well cell culture plates at the initial density of 5 × 10^5^ cells/ml in Dulbecco’s modified Eagle medium (Hyclone, Massachusetts, USA) containing 10% fetal bovine serum and 1% penicillin and streptomycin. The cells were cultured in a 5% CO_2_ incubator at 37°C.

### NAD^+^ assay

As previously described (19), NAD^+^ concentrations were determined by the recycling assay. Briefly, samples were extracted in 0.5 N perchloric acid. After centrifugation at 12,000 g for 5 min, the supernatant was obtained, which was neutralized to pH 7.2 by using 3 N potassium hydroxide and 1 M potassium phosphate buffer. After centrifugation at 12,000 g for 5 min, the supernatants were mixed with a reaction media containing 1.7 mg 3-[4,5-dimethylthiazol-2-yl]-2,5-diphenyl-tetrazolium bromide (MTT), 1.3 mg alcohol dehydrogenase, 488.4 mg nicotinamide, 10.6 mg phenazine methosulfate, and 2.4 mL ethanol in 37.6 mL Gly-Gly buffer (65 mM, pH 7.4). After 10 min, the A560nm was determined by a plate reader, and the readings were calibrated with NAD^+^ standards. The protein concentrations were assessed by BCA Protein Assay (Thermo Scientific,Waltham,MA,USA).

### Flow cytometry-based Annexin V/7-AAD assay

The apoptosis level of PC12 cells was determined by ApoScreen Annexin V kit (SouthernBiotech, Birmingham, AL, USA) according to the manufacturer’s protocol. Briefly, cells were harvested by 0.25% trypsin solution, then washed by cold PBS. The cells were resuspended in 100μl cold 1X binding buffer (10 mM HEPES, pH 7.4, 140 mM NaCl, 2.5 mM CaCl_2_, 0.1% BSA). Then 5 μL of labeled Annexin V was added into the cell suspension. After incubation for 15 min on ice, the solution contained 200 μL 1X binding buffer and 5 μL 7-AAD was added into the cell suspensions. The number of stained cells was measured by a flow cytometer (FACSAria II, BD Biosciences).

### ATP assay

Intracellular ATP levels were determined by an ATP Bioluminescence Assay Kit (Roche Applied Science, Mannheim, Germany), according to the manufacturer’s protocols. Briefly, PC12 cells were washed once with PBS, lysed using the cell lysis reagent, and then the cell samples were mixed with 50 μl of the luciferase reagent. The chemiluminescence of the samples was detected using a plate reader (Synergy2; Biotek, Winooski, Vermont, USA). The ATP levels were normalized to the protein concentrations of the samples, which were determined by BCA Protein Assay (Thermo Scientific, Waltham, MA, USA).

### Real-Time PCR assay

After total RNA was extracted (TaKaRa MiniBEST Universal RNA Extraction Kit, Takara Bio, Dalian, China) from PC12 cells, 500 ng total RNA was reverse-transcribed to cDNA (Prime-Script RT reagent kit, Takara Bio, Dalian, China). The parameters set for reverse transcription were as follows: 37°C for 15 min, then 85°C for 15 s. Quantitative RT-PCR was performed by using SYBR Premix Ex Taq (Takara Bio, Dalian, China) and the following primers: Nampt (sense 5’-TATTCTGTTCCAGCGGCAGA -3’ and anti-sense 5’-GACCACAGACACAGGCACTGA -3’); GAPDH (sense 5’-CCTGCACCACCAACTGCTTA -3’ and anti-sense 5’-GGCCATCCACAGTCTTCTGA -3’). Assays were performed according to the following procedure: Denatured at 95°C for 10 s, followed by 40 cycles of 95°C for 5 s and 60°C for 30 s. The data were analyzed by using the comparative threshold cycle method, and the results were expressed as fold differences normalized to the mRNA level of GAPDH.

### Western blot assay

After the various treatments, PC12 cells were washed once with PBS. Then the PC12 cells were harvested and lysed in RIPA buffer (Millipore, Temecula, California, USA) containing 1% protease inhibitor cocktail (CWBio, Beijing, China) and 1 mM phenylmethanesulfonyl fluoride. Lysates were centrifuged at 12,000 rpm for 10 min at 4°C. After quantifications of the protein samples using BCA Protein Assay Kit (Pierce Biotechnology, Rockford, Illinois, USA), 30 μg of total protein was electrophoresed through a 10% sodium dodecyl sulfate–polyacrylamide gel and then transferred to 0.45-μm nitrocellulose membranes. The blots were incubated overnight at 4°C with rabbit polyclonal Nampt antibody (1:2000 dilution, Abcam, Cambridge, UK) in TBST containing 1% bovine serum albumin (BSA) and then incubated with horse radish peroxidase-conjugated secondary antibody (1:3000 dilution, Epitomics, Hangzhou, Zhejiang Province, China) in TBST containing 1% BSA at room temperature for 1 hr. A GAPDH antibody (proteintech, Wuhan, China) was used to normalize the sample. The intensities of the bands were quantified by densitometry using Gel-Pro Analyzer (Media Cybernetics, Silver Spring, Maryland, USA).

### Intracellular lactate dehydrogenase (LDH) assay

Cell death was determined by measuring the intracellular lactate dehydrogenase (LDH) activity of the cells. In brief, cells were lysed for 20 min in lysing buffer containing 0.04% Triton X-100, 2 mM HEPES and 0.01% bovine serum albumin (pH 7.5). Then 50 μL cell lysates were mixed with 150 μL reaction buffer containing 0.34 mM NADH and 2.5 mM sodium pyruvate (PH 7.5). The A_340nm_ changes of the samples were monitored over 90 s. Percentage of cell death was calculated by normalizing the LDH values of samples to the LDH values of the lysates of control.

### RNA interference

When PC12 cells were approximately 40% confluent, the cells were transfected with Nampt siRNA sequences (sense 5’-AGUAAGGAAGGUGAAAUACTT-3’ and antisense 5’-GUAUUUCACCUUCCUUACUTT-3’) and the scrambled control siRNA sequences. Lipofectamine 2000 (Invitrogen, Carlsbad, California, USA) was used for transfection according to the manufacturer’s protocol. For each well of a 24-well plate, 100 μl Opti-MEM containing 0.06 nmol of the siRNA sequences and 2.5 μl lipofectamine 2000 was added into 500 μl culture media of the cells. After 6 hrs, the media was replaced by DMEM containing 10% fetal bovine serum.

### Statistical analyses

All data are presented as mean ± SEM. Data were assessed by one-way ANOVA, followed by Student – Newman – Keuls *post hoc* test, except where noted. *P* values less than 0.05 were considered statistically significant.

## Results

### 1. Nampt plays siginificant roles in both the NAD^+^ synthesis and survival of PC12 cells under basal conditions

To investigate the roles of Nampt in the NAD^+^ metabolism of PC12 cells under basal conditions, we determined the effects of the Nampt inhibitor FK866 on the intracellular NAD^+^ levels of the cells. FK866 at the concentrations of 2, 5, 10, 20, and 50 nM led to profound decreases in the intracellular NAD^+^ levels (Fig. 1A), indicating a critical role of Nampt in the NAD^+^ synthesis in the cells under basal conditions. We also applied FACS-based Annexin V/7-ADD assay to determine the roles of Nampt in the basal survival of the cells, showing that 10 nM FK866 led to a significant increase in early-stage apoptosis, while both 20 nM and 50 nM FK866 led to significant increases in early-stage apoptosis (Annexin V^+^/7-ADD^−^), late-stage apoptosis (Annexin V^+^/7-ADD^+^) and necrosis (Annexin V^−^/7-ADD^+^) (Figs. 1B and 1C). FK866 dose-dependent increased cell death, which equlas to the sum of early-stage apoptosis (Annexin V^+^/7-ADD^−^), late-stage apoptosis (Annexin V^+^/7-ADD^+^) and necrosis (Annexin V^−^/7-ADD^+^) (Figs. 1B and 1C).

**Fig. 1.**
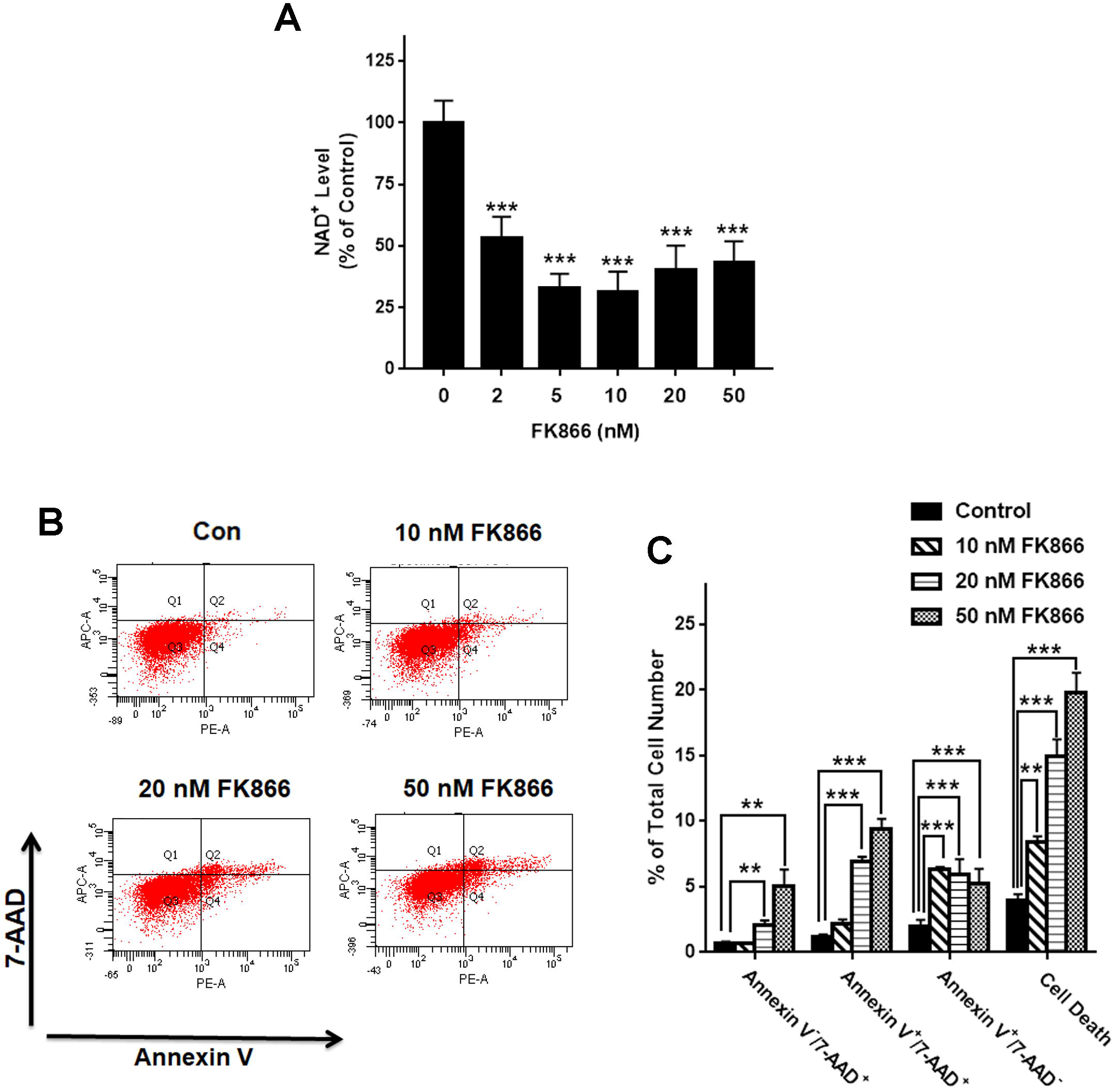
Nampt plays a siginificant role in the NAD^+^ synthesis and basal survival of differentiated PC12 cells. (A) FK866 at the concentrations of 2, 5, 10, 20, and 50 nM led to profound decreases in the intracellular NAD^+^ levels. NAD^+^ assays were conducted 12 hrs after the cells were treated with 2, 5, 10, 20, and 50 nM FK866. *N* = 8. The data were pooled from three independent experiments. ***, *P* < 0.001. (B,C) 10 nM FK866 led to a significant increase in early-stage apoptosis, while both 20 nM and 50 nM FK866 led to significant increases in early-stage apoptosis (Annexin V^+^/7-ADD^−^), late-stage apoptosis (Annexin V^+^/7-ADD^+^) and necrosis (Annexin V^−^/7-ADD^+^). PC12 cells were treated with 2, 5, 10, 20, and 50 nM FK866 for 24 hrs. *N* = 6. The data were pooled from three independent experiments. **, *P* < 0.01; ***, *P* < 0.001.

We also applied Nampt siRNA to determine the effects of decreased Nampt on the survival of PC12 cells. Treatment of the cells with Nampt siRNA led to a significant decrease in the Nampt level (Supplemental Fig. 1A). The reduction of Nampt led to both a significant increase in the cell death (Supplemental Fig. 1B) and a significant decrease in the cell survival (Supplemental Fig. 1C).

### 2. H_2_O_2_ produced significant decreases in both mRNA and protein levels of Nampt in PC12 cells

We determined the effects of H_2_O_2_ on the Nampt mRNA levels of the PC12 cells at 12 or 20 hrs after the H_2_O_2_ treatment. Treatment of the cells with 0.1 or 0.3 mM H_2_O_2_ led to significant increases in the Nampt mRNA level of the cells at 12 hrs after the H_2_O_2_ treatment (Fig. 2A). In contrast, H_2_O_2_ led to significant decreases in the Nampt mRNA level at 20 hrs after the H_2_O_2_ treatment (Fig. 2A).

**Fig. 2.**
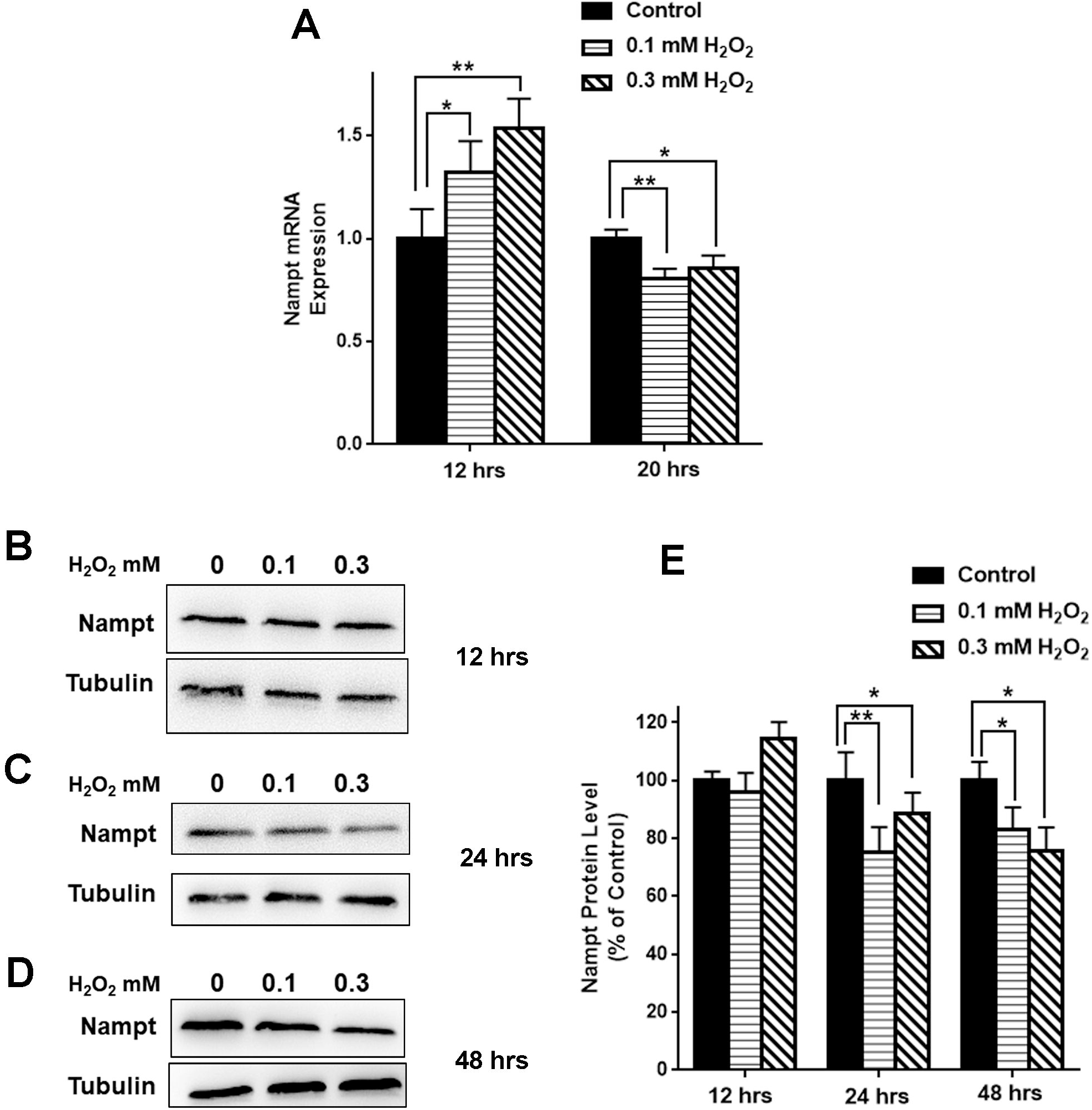
H_2_O_2_ produced significant decreases in both Nampt mRNA levels and Nampt protein levels of PC12 cells. (A) Treatment of PC12 cells with 0.1 mM or 0.3 mM H_2_O_2_ led to significant increases in the Nampt mRNA level of the cells at 12 hrs after the H_2_O_2_ treatment. In contrast, H_2_O_2_ led to significant decreases in the Nampt mRNA level of the cells at 20 hrs after the H_2_O_2_ treatment. PC12 cells were treated with 0.1, 0.3 mM H_2_O_2_ for 12 or 20 hrs. *N* = 6. The data were pooled from three independent experiments. *, *P* < 0.05; **, *P* < 0.01. (B) Effects of H_2_O_2_ on the protein level of Nampt at 12 hrs after the H_2_O_2_ treatment. (C) Effects of H_2_O_2_ on the protein level of Nampt at 24 hrs after the H_2_O_2_ treatment. (D) Effects of H_2_O_2_ on the protein level of Nampt at 48 hrs after the H_2_O_2_ treatment. (E) H_2_O_2_ did not significantly affect the level of Nampt at 12 hrs after the H_2_O_2_ treatment. In contrast, H_2_O_2_ led to significant decreases in the Nampt protein level of the cells at both 24 and 48 hrs after the H_2_O_2_ treatment. PC12 cells were treated with 0.1, 0.3 mM H_2_O_2_ for 12, 24, or 48 hrs. *N* = 9. The data were pooled from three independent experiments. *, *P* < 0.05; **, *P* < 0.01.

We also determined the effects of H_2_O_2_ on the Nampt protein levels of the cells at 12, 24 or 48 hrs after the H_2_O_2_ treatment. H_2_O_2_ did not significantly affect the level of Nampt at 12 hrs after the H_2_O_2_ treatment (Figs. 2B and 2E). In contrast, H_2_O_2_ led to significant decreases in the Nampt protein level at both 24 (Figs. 2C and 2E) and 48 hrs (Figs. 2D and 2E) after the H_2_O_2_ treatment.

### 3. H_2_O_2_-produced decrease in the Nampt levels plays a significant role in H_2_O_2_-produced cell death

To investigate the biological consequences of the H_2_O_2_-produced decrease in the Nampt levels of PC12 cells, we applied FACS-based Annexin V/7-ADD assay to determine the effects of the Nampt inhibitor FK866 on H_2_O_2_-produced cell death. FK866 significantly exacerbated H_2_O_2_-produced early-stage apoptosis (Annexin V^+^/7-ADD^−^), late-stage apoptosis (Annexin V^+^/7-ADD^+^) and necrosis (Annexin V^−^/7-ADD^+^) (Figs. 3A and 3B). Nampt siRNA treatment also significantly exacerbated the H_2_O_2_-produced cell death, asssesed by LDH assay (Supplemental Fig. 1D). To further investigate the biological consequences of the H_2_O_2_-produced decreases in the Nampt protein levels of PC12 cells, we determined the effects of the Nampt activator P7C3 on the H_2_O_2_-produced cell death. P7C3 significantly attenuated H_2_O_2_-produced decrease in the cell survival (Fig. 3C). Our study also showed that FK866 and H_2_O_2_ produced synergistic effects to decrease the intracellular ATP levels (Fig. 4).

**Fig. 3.**
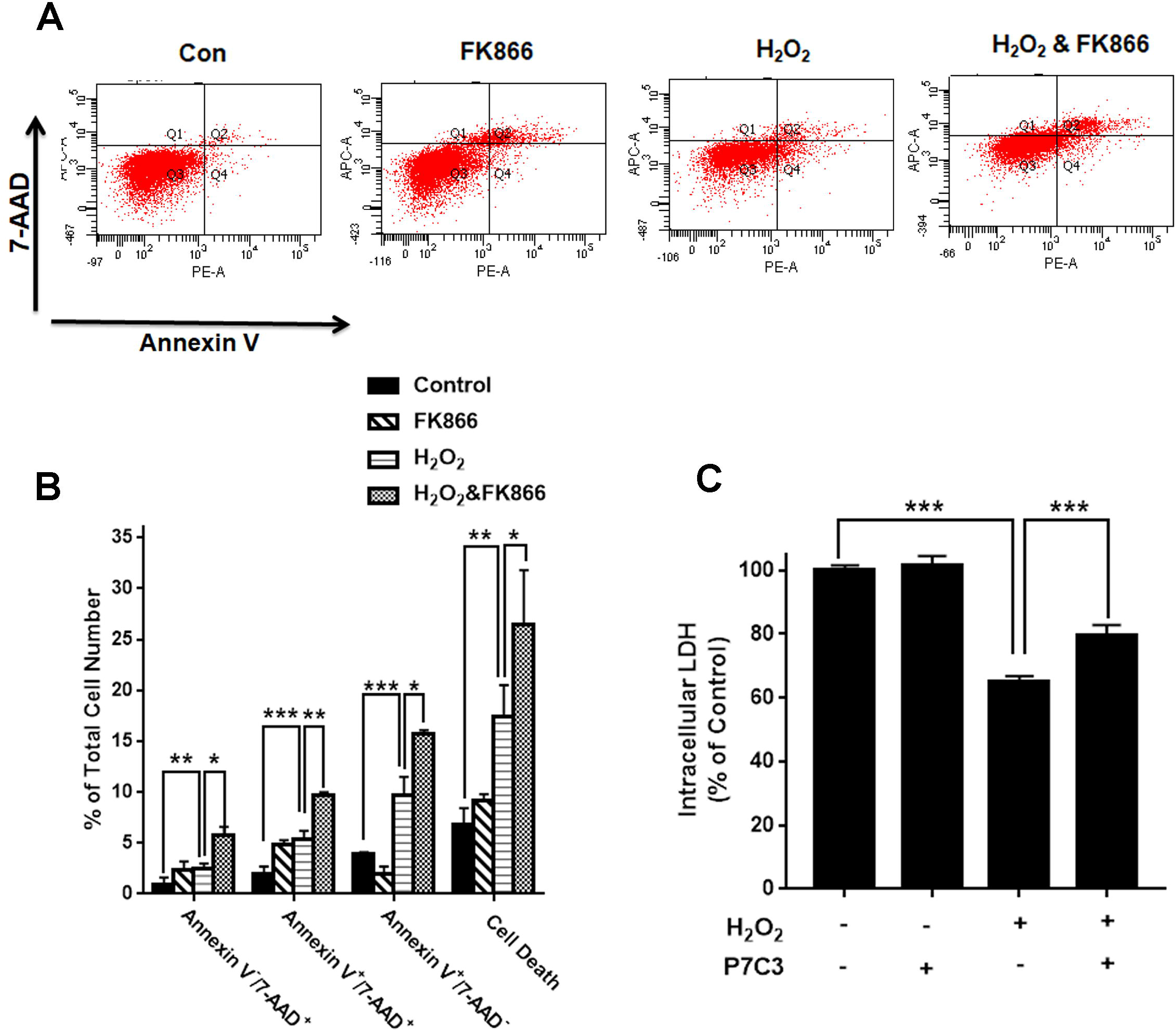
Nampt inhibitor FK866 and Nampt activator P7C3 significantly exacerbated or attenuated H_2_O_2_-produced cell death of differentiated PC12 cells, respectively. (A, B) FK866 significantly exacerbated H_2_O_2_-produced early-stage apoptosis (Annexin V^+^/7-ADD^−^), late-stage apoptosis (Annexin V^+^/7-ADD^+^) and necrosis (Annexin V^−^/7-ADD^+^) of the cells. PC12 cells were pre-treated with 10 nM FK866 for 30 min, and then co-treated with 0.3 mM H_2_O_2_ for 23.5 hrs. *N* = 6. The data were pooled from three independent experiments. *, *P* < 0.05; **, *P* < 0.01; ***, *P* < 0.001. (C) Nampt activator P7C3 significantly attenuated H_2_O_2_-produced decrease in the cell survival. PC12 cells were pre-treated with 10 μM P7C3 for 4 hrs, and then co-treated with 0.3 mM H_2_O_2_ for 20 hrs. *N* = 9. The data were pooled from three independent experiments. ***, *P* < 0.001.

**Fig. 4.**
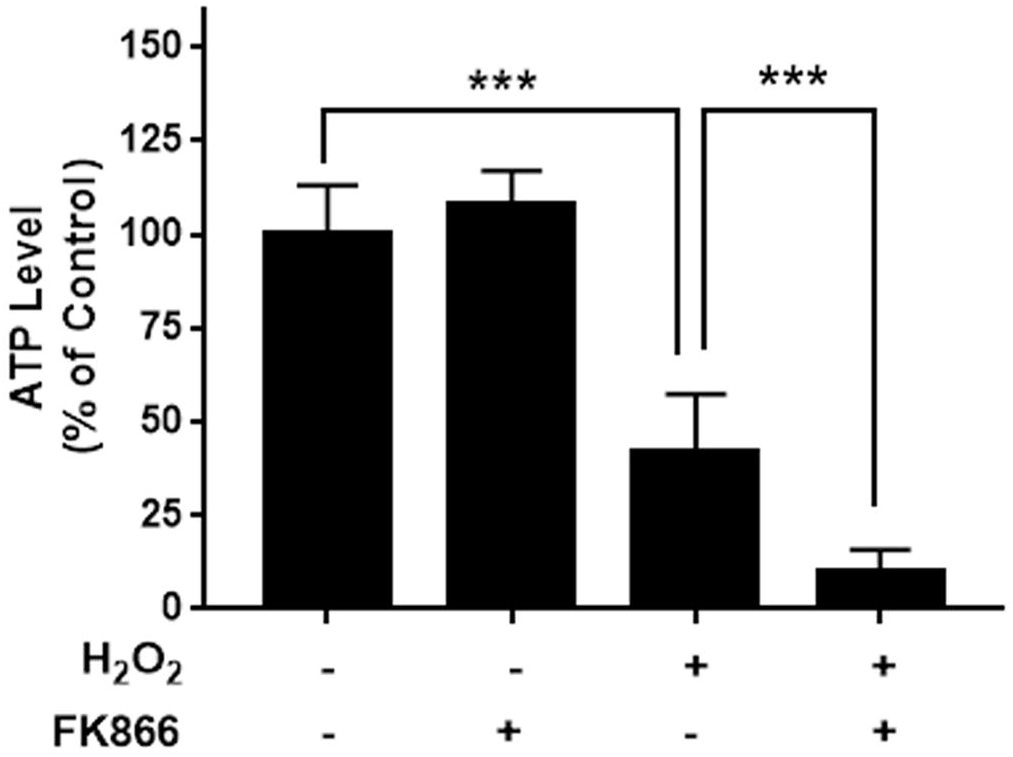
Nampt inhibitor FK866 significantly exacerbated H_2_O_2_-produced decreases in the intracellular ATP levels of PC12 cells. FK866 and H_2_O_2_ produced synergistic effects to decrease the intracellular ATP levels. PC12 cells were pre-treated with 10 nM FK866 for 30 min, and then co-treated with 0.3 mM H_2_O_2_ for 1 hr. *N* = 6. The data were pooled from three independent experiments. ***, *P* < 0.001.

## Discussion

The major findings of our current study include: First, Nampt plays significant roles in both the NAD^+^ synthesis and survival of the cells under basal conditions; second, H_2_O_2_ produced significant decreases in both the mRNA and the protein levels of Nampt; and third, H_2_O_2_ induced cell death partially by producing the decreases in the mRNA and protein levels of Nampt, since the Nampt inhibitor or the Nampt activator significantly exacerbated or attenuated the H_2_O_2_-induced cell death, respectively.

Cumulating evidence has indicated critical roles of impaired NAD^+^ metabolism in both aging and a number of major diseases (1, 2, 3), indicating critical significance to investigate the mechanisms underlying NAD^+^ deficiency under these conditions (1, 2, 3). Under in the tissues or organs certain pathological conditions, including the metabolic organs that are exposed to HFD (16), Drosophila pink1 mutants (17), and aging (16, 18), decreased levels of Nampt have been reported. However, there is significant deficiency in the mechanisms underlying the Nampt alterations under pathological conditions.

Our current study has indicated that H_2_O_2_ was capable of decreasing both the mRNA and protein levels of Nampt in PC12 cells. Since there is significantly increased oxidative stress in aging and the pathological conditions (20–22), our finding has suggested a novel mechanism underlying the decreased Nampt in aging and multiple pathological conditions. Due to the critical roles of NAD^+^ deficiency in aging and multiple major diseases, our study has suggested a significant mechanism underlying the pathological alterations of in aging and the diseases: The oxidative stress in the diseases and aging can lead to decreased Nampt, leading to decreased NAD^+^ synthesis thus resulting multiple pathological changes.

Our study has also indicated that Nampt plays a significant role in oxidative stress-induced cell death, since the Nampt inhibitor or the Nampt activator significantly exacerbated or attenuated the H_2_O_2_-induced cell death, respectively. These observations have suggested a novel mechanism underlying oxidative stress-induced cell death at least for some cell types.

There are two NAD^+^ synthesis pathways in cells, while the roles of Nampt in the both general NAD^+^-generating capacity and survival of various cell types under basal conditions are largely unclear. Our current study has indicated that Nampt plays critical roles in both the NAD^+^ synthesis and cell survival of differentiated PC12 cells under basal conditions, since the Nampt inhibitor FK866 produced profound decreases in both the intracellular NAD^+^ levels and survival of the cells under basal conditions.

Collectively, our study has indicated significant roles of Nampt in the survival of PC12 cells under both basal and oxidative stress conditions. Our study has also indicated a novel mechanism underlying the decreases in Nampt in multiple pathological conditions and aging. Moreover, our study has suggested a novel mechanism underlying oxidative cell death.

## Acknowledgment

The authors would like to acknowledge the financial support by two research grants from a Major Special Program Grant of Shanghai Municipality (Grant # 2017SHZDZX01) (to W.Y.) and a Major Research Grant from the Scientific Committee of Shanghai Municipality #16JC1400502 (to W.Y.).

## Legends of Supplemental Figures

**Supplemental Figure 1. Nampt siRNA-produced reduction of Nampt led to a significant increase in differentiated PC 12 cell death.** (A) Treatment of the cells with Nampt siRNA led to a significant decrease in the Nampt level. PC12 cells were transfected with control or Nampt siRNA sequences for 48 hrs. *N* = 12. The data were pooled from three independent experiments. ***, *P* < 0.001. (B, C) The reduction of Nampt led to a significant increase in the cell death. PC12 cells were transfected with control or Nampt siRNA sequences for 48 hrs. *N* = 9. The data were pooled from three independent experiments. *, *P* < 0.05; ***, *P* < 0.001. (D) The reduction of Nampt exacerbated H_2_O_2_-produced cell death. PC12 cells were transfected with control or Nampt siRNA sequences for 24 hrs, and then incubated by 0.3 mM H_2_O_2_ for 24 hrs. *N* = 9. The data were pooled from three independent experiments. ***, *P* < 0.001.

