## Supplementary figures and images for "Oxidative stress induces cell death partially by decreasing both mRNA and protein levels of nicotinamide phosphoribosyltransferase in PC12 cells"

### Supplemental Fig. 1

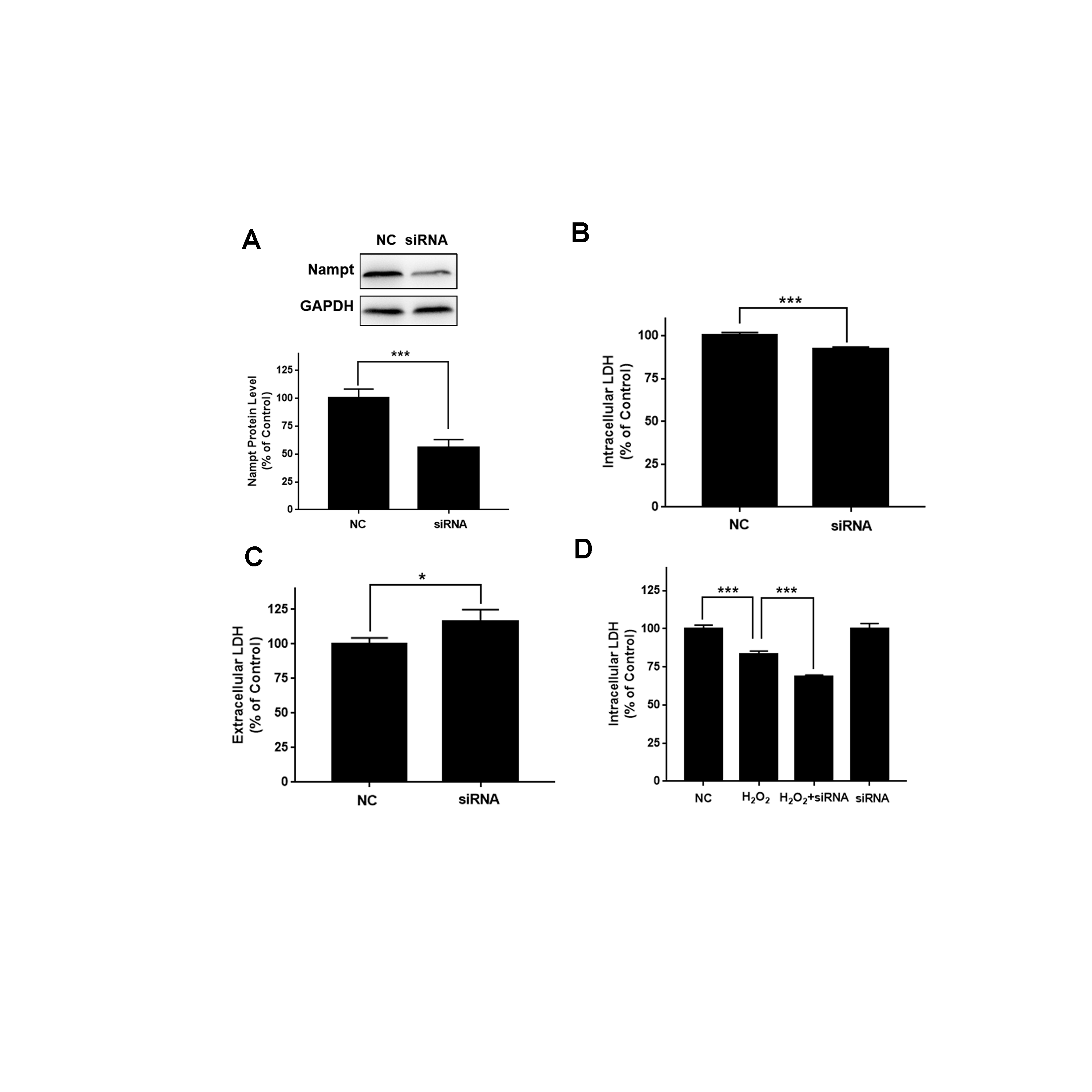
